# Bashing irreproducibility with *shournal*

**DOI:** 10.1101/2020.08.03.232843

**Authors:** Tycho Kirchner, Konstantin Riege, Steve Hoffmann

## Abstract

The Linux shell is arguably one of the most important computational tools across various scientific disciplines. Its high flexibility makes it the platform of choice for many file operations and smaller scripting tasks. Also, many stand-alone programs are called from the Linux shell - typically followed by multiple command-line parameters. However, in larger analysis projects, keeping track of the work quickly becomes challenging, as a typical shell workflow involves the iterative execution of commands with many parameters, modification of scripts, and editing of configuration files. Too often, researchers find themselves in the uncomfortable position that a computational result generated a few weeks ago can no longer be reproduced, despite having taken great care documenting the work manually. On the other hand, there is a lack of tools able to record the researcher’s shell activity automatically with reasonably low runtime- and storage overhead. To close this critical gap, we developed *shournal*, a program that tightly integrates with the Linux shell and automatically records every shell command along with the files it reads or writes. Besides logging command- and file metadata, such as working directory, file path, and checksums, *shournal* can be configured to archive scripts or configuration files that are not regularly under version control via *git* or *svn*.

## 2 Main

*shournal* provides a command-line interface to allow fast user queries. For instance, for a given file, the commands that created or used it can be retrieved (Fig. 1c). Automatically archived scripts or configuration files are restorable, enabling a re-execution of commands, even if the original files have been modified or deleted. Saved file checksums help to quickly determine which input files or scripts changed. Other options include queries for files modified during a given period, the command history of a project directory, or commands executed during a specific shell session.

**Fig. 1:**
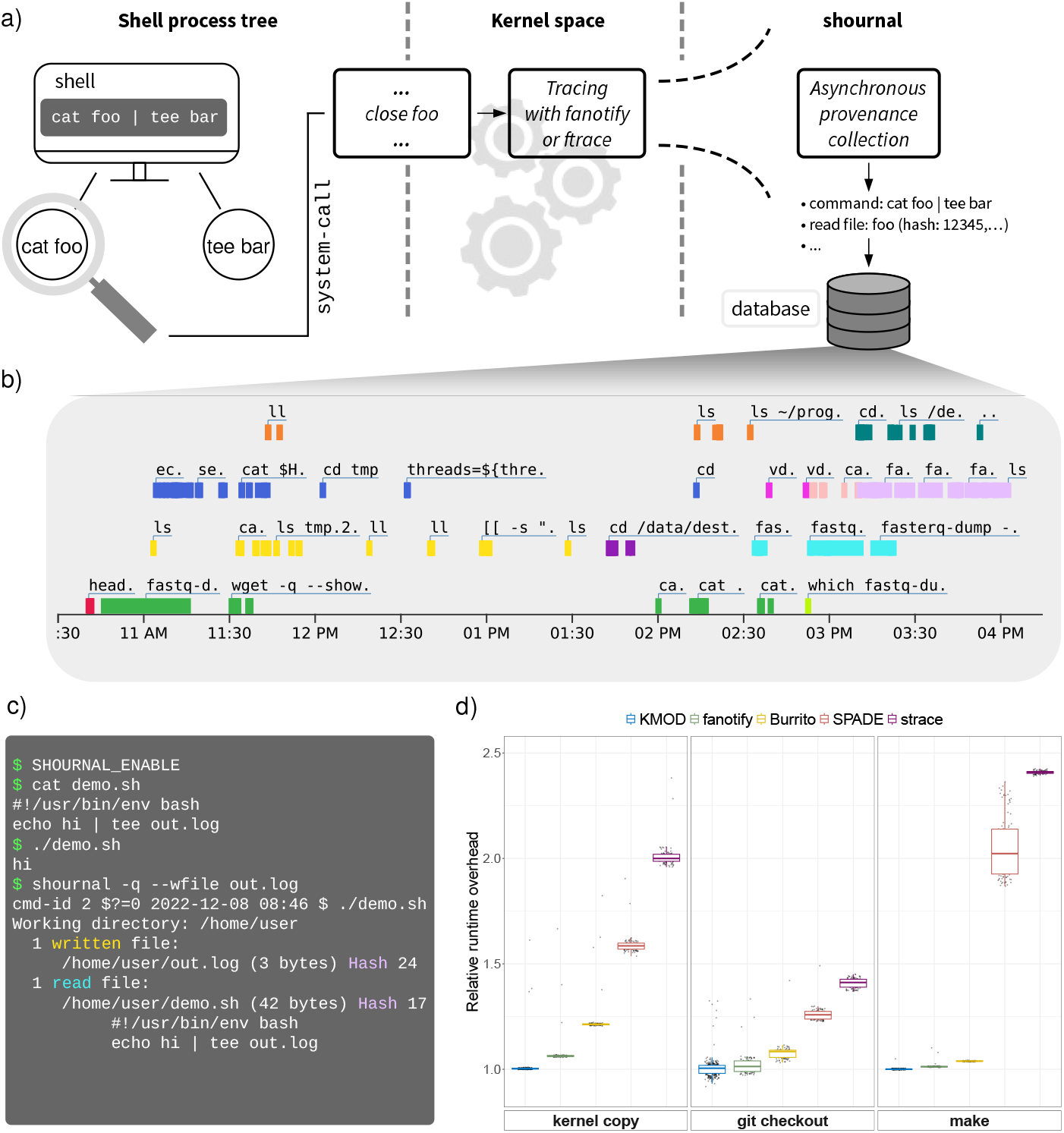
Record shell commands and used files with shournal. (a) Schematic illustration of a shell session observed by *shournal*. The whole process tree of the command `cat foo` | `tee bar` is monitored for file close system calls. When `foo` is closed the event is traced in kernel space within process context while provenance collection runs asynchronously. Finally metadata and checksums of the used files are stored alongside the corresponding shell command within *shournal*’s database. (b) Visualization of the shell command history. *shournal* exports the command history into an interactive HTML-plot. Commands, which were executed within a given shell session, are marked with the same color. Parallel shell sessions are vertically stacked. (c) An example shell session observed by *shournal*. The executed script `demo.sh` creates a file `out.log`. Next, *shournal*’s database is queried for commands which created or modified `out.log` using its `--wfile` option. *shournal* reports the command, file checksums and the archived script. (d) *shournal*’s tracing performance in various scenarios as relative runtime overhead. Boxes for both, kernel module- (KMOD) and fanotify backend are displayed. For comparison, our measured tracing overhead of *Burrito*, SPADE and the *ptrace*-based *strace* is shown as well.

Such queries may be helpful for virtually all scientists working on the shell, especially when applications with various input options are used. An example from computational chemistry is the deep learning framework *OpenChem* (Korshunova, 2021). Parameters of the deep learning models, like the number of epochs, are defined in configuration files. However, values can be overridden via the command line. *shournal* tracks both in conjunction, making later re-execution easier. In addition to the console output, *shournal* generates an interactive graphical map of commands (Fig. 1b). Clicking on a command gives supplementary information, e.g., time of execution, or archived scripts.

In recent years, many tools addressing the general problem of computational reproducibility have been developed (Ruiz, Richard; Ivie and Thain, 2018) – often geared toward the reproduction of pipelines across platforms to share them with other researchers. Prominent examples include *CARE* (Janin, Vincent), *CDE* (Guo, 2012), and *ReproZip* (Chirigati et al., 2016). Using the Linux’s *ptrace*, these tools are able to automatically create an executable, easy-to-share archive of an entire pipeline and its dependencies. However, due to the necessarily high runtime- (Fig. 1d) and storage overhead, a continuous usage for all shell commands is not generally advisable. We therefore assume that such archives are, in practice, created at a project’s essential milestones, requiring all intermediate steps to be reproducible. As outlined above, this can be quite challenging, especially when several days or weeks have passed until those archivers run. *shournal* promotes the usage of these tools by facilitating the reconstruction of the processing steps performed before the fully reproducible experiment is created.

Workflow engines like *Snakemake* (Köster and Rahmann, 2012) or *Nextflow* (Di Tommaso, 2017) provide their script language to describe pipelines, also aiming at enhancing computational reproducibility. However, analysis projects are not always consequently carried out within these frameworks. *shournal* can simplify the creation of such workflows by sharing its data using JSON. Due to the low-level nature of the data, observed shell commands can often be directly transformed into *Snakemake* rules utilizing the software at https://github.com/snakemake/shournal-to-snakemake. A rule’s input- and output section is generated from the captured file events.

While developing *shournal*, we reviewed other tracers, like *record-and-replay systems*, SPADE (Gehani and Tariq, 2012), or *Burrito* (Guo and Seltzer, 2012). However, the conceptually more extensive design goals of these tools result in high runtime- and storage overheads and a rather difficult use (see Supplementary Material). *shournal* is designed to run permanently with low overhead, even when tracing demanding pipelines. Typical workflows yield a runtime overhead of less than 0.5% (Fig. 1d) while the storage overhead is small, i.e., a few megabytes in case of tens of thousands of file events. Over-heads are kept low using three key concepts: First, tracing is implemented using *ftrace* and *tracepoints* from *shournal’s* own kernel module (KMOD) or the Linux kernel’s *fanotify* filesystem API, instead of, e.g., the much slower *ptrace* (Gebai and Dagenais, 2018). Second, tracing of file actions is limited to the comparatively rare close operation and lets the traced process return quickly by delegating further provenance collection to another thread (Fig. 1a). Third, by default, *shournal* primarily captures file metadata such as path, size, modification-time, and (partial) checksums, reducing the data read or written to. More details, performance considerations, and limitations are discussed in the Supplementary Material.

*shournal* allows to record, search and visualize the work carried out on the shell with configurable low runtime- and storage overhead. Therefore, it can play an essential role in facilitating the reproducibility of analyses by allowing to reconstruct, summarize, and to resume projects quickly. With the *shournal-to-snakemake* converter, we demonstrate that the collected data may be used as a basis for the creation of workflows. By narrowing the gap between poorly documented ad-hoc work on the shell and the generation of sophisticated workflows or fully reproducible experiments using *CARE, CDE* or *ReproZip, shournal* can help to save time and to improve scientific practice.

## Availability

*shournal* is available at https://github.com/tycho-kirchner/ shournal under GNU General Public License v3.0.

