## Supplementary File for "Bashing irreproducibility with *shournal*"

### Bashing irreproducibility with *shournal* - Supplementary material

#### 1 Related software

In this study, we reviewed computational reproducibility tools concerning their suitability for permanently tracing the user's shell commands. One of the most comprehensive approaches is taken by so-called *record-and-replay* systems like *Arnold* (Devecsery et al., 2014) or *rr* (Record-Replay) (O'Callahan et al., 2017), which aim at recording entire process trees with plenty of details, e.g., used system calls and their return value, to allow "replaying" them at a later time. However, while being very powerful and suitable for, e.g., debugging, forensics, or finding data races, the amount of necessary tracing and logging to replay processes may be too high when computationally demanding pipelines are monitored regularly. For instance, *Arnold* reports a runtime overhead of up to 100% for a simple `cvs checkout` and 1 TB of disk space per year and workstation. Further, to use *Arnold*, a custom Linux kernel must be compiled, constituting a high burden for administrators or users. The more recent *rr* is based on an optimized *ptrace* using *seccomp-bpf* filters and does not require a custom kernel. However, it reports an almost eightfold runtime overhead for the example of compiling source code using *make*.

SPADE (Gehani and Tariq, 2012) allows recording provenance on the system level using the Linux kernel's built-in *auditd* infrastructure. While it does not record shell commands, in principle, a plugin could add such functionality. However, while the authors report a runtime overhead of about 10% when monitoring a web server or a genomic sequence alignment, we find overheads partially exceeding 100% in our own benchmark, discouraging us from using SPADE as a tracing backend.

The bash plugin of *Burrito* (Guo and Seltzer, 2012) associates shell commands and their used files based on tracing with *SystemTap* (SystemTap, 2023). However, it yields a runtime overhead of partially more than 20% in our benchmark, which is, as *shournal* demonstrates, substantially higher than necessary. Further, *Burrito* relies on the NILFS versioning filesystem (Konishi et al., 2006) to save previous file versions posing a severe limitation for institutions built around network filesystems such as NFS. Finally, *Burrito*'s unconditional preservation of all file versions could quickly result in unacceptable storage overhead.

### 2 Supplementary Methods

##### 2.0.1 Tracing

*shournal* implements its design goal to associate shell commands and their used files through kernel space tracing. Specifically, *shournal* installs a global hook within the Linux kernel, which runs whenever any process on the system closes the last instance of a file descriptor. Within that hook, it is determined if the executing process is part of an observed shell command, whereupon a raw reference on the file path (a kernel object, not a string) is taken and buffered. This allows the monitored process to resume quickly while further processing steps run asynchronously in another thread. *shournal* provides two backends to trace file close-events of specific processes: its own kernel module *shournalk* and another, *fanotify*-based backend.

*shournalk* is based on tracepoints and the *ftrace* framework, allowing to run custom code at specific execution paths without kernel recompilation (Gebai and Dagenais, 2018). Only three events are traced. In addition to tracing the closing of files, the *exit* of processes and *fork* events are kept track of. Tracing of the latter allows to maintain a list of shell processes and their descendants. A process can initially be marked for observation using *shournalk*'s sysfs-interface (Kerrisk, 2010, chap. 14.1).

The *fanotify*-based backend employs the kernel-native *fanotify* filesystem API to register for file close-events. However, while *fanotify* allows event subscription for whole mount points (Kerrisk, 2010, chap. 14.7), it provides no direct way to monitor only file events of specific processes. To remedy this shortcoming, for each shell command, *shournal* creates a unique unshared mount namespace (Kerrisk, 2010, chap. 28.2.1) and ensures that file operations of the shell and its child processes refer to it.

The *fanotify*-backend provides a shared library, hooked into the shell's process and a setuid (*suid*) program (Kerrisk, 2010, chap. 9.3) performing the privileged actions of unsharing and joining mount namespaces as well as marking mount points with *fanotify*. Specifically, the shared library masks the library calls *open* and *execve*. The former ensures that files within the parent shell and its subshells are opened relative to the new mount namespace, while the latter redirects to *shournal*'s *suid* binary which performs the privileged *setns* syscall to join the new mount namespace, executing the original program afterward. The mount-namespace is, by default, inherited during *fork*, notable exceptions being, e.g., container solutions like *Docker*. As a result, applications unsharing the mount namespace themselves cannot be traced from an outer layer. However, *shournal* provides a dedicated *Docker* edition which does allow a shell running within *Docker* to be observed.

While *shournalk* is faster (see section Performance), has fewer limitations, and alters the user-space environment to a lesser extent, the *fanotify* backend may be of particular interest to institutions where the installation of foreign kernel modules is discouraged.

##### 2.0.2 Provenance collection

An asynchronous provenance collection thread consumes the file close events, which were buffered for observed shell processes and their children. It filters files according to the user's preferences, e.g., by archiving only scripts ending with *.sh* or ignoring events from the system's temporary directory. To ensure the identity of a given file, beyond metadata like name, size, and modification time, a checksum of the file content is calculated using *xxHash* https://github.com/Cyan4973/xxHash. As hashing large files in their entirety could introduce a considerable slowdown, only $N$ chunks of length $b$ bytes of a file are digested at regular offsets calculated from the file size $s$. Starting from the beginning of a file, the seek-step $p$ is $p = \lfloor \frac{s}{N} \rfloor$ bytes. Small files, where $p \leq b$, are hashed completely. Three chunks of 256 file bytes are digested, by default, keeping the overhead low. On the downside, files of equal size with the same content in the hashed regions but different content in others are falsely reported as equal. While we did not encounter such an error in practice, *shournal*'s hash settings remain configurable to digest more parts of a file, if appropriate. Obtaining the correct checksum of a file's "final" version is guaranteed to be free from race conditions as long as all writers are part of the observed process tree: at some point, the last writer closes the file, after which the final hashing is performed. Size, checksum, and optionally modification time allow for later file provenance queries with high accuracy independent of the filename. Thus, *shournal* abstains from tracing rename operations.

In the first instance, to be fast and lightweight, metadata and archived scripts are stored in a binary, partially compressed file and later, without time pressure, finally stored in a sqlite database by a low-priority background daemon. The maximum number of archived scripts and file events per command is configurable, so, by default, a backup script cannot flood *shournal*'s database. To further reduce disk usage, files are archived in a deduplicated manner.

##### 2.0.3 Performance

To analyze *shournal*'s performance, we designed a benchmark reflecting scenarios with intensive file access. During *kernel copy*, for instance, more than 120 000 files were read and written within a few seconds. In addition, we benchmarked a *git checkout* of the Linux kernel source code and the compilation of *elfutils* with *make*. For each command, the time of a "cold cache" run was measured at least 100 times, while between each run the file cache of the Operating system (OS), as well as the disk cache, was cleared.

We find that the median runtime overhead is less than 0.5% for the kernel module backend and less than 6.3% for the *fanotify* backend in all examined cases. These results demonstrate the applicability of our tool.

For comparison with *Burrito*, we benchmarked its *SystemTap*-script (Guo and Seltzer, 2012). The runtime overhead of partially exceeding 20% already occurs due to plain tracing and event-logging. No further processing or file-versioning using the NILFS filesystem was involved, so we consider this the lower bound of the performance penalty one has to expect when using this tool.

For comparison with SPADE, we installed and configured its *auditd* reporter ( `add reporter Audit`). The runtime overhead of partially exceeding 100% already occurs due to plain tracing and event processing (without disk logging), so we conclude that the runtime overhead might be even higher in practice. Note that we did not measure SPADE's *camflow* backend due to our initial requirement of not depending on a self-compiled kernel.

Finally, as a representative for *ptrace*-based tools, we measured runtimes under the tool *strace* (Kerrisk, 2010, appx. A) and found large overheads of partially exceeding 140%.

The storage overhead of *shournal*'s database critically depends on the user configuration. Using defaults, the `cp`-benchmark yields a disk usage of 174 bytes per file event. Thus, one GiB of disk space is sufficient for recording approx. 6 million events. For a regular shell user, we estimate a storage requirement of less than 100MiB per month. Furthermore, *shournal* provides a rich command line interface to delete unneeded entries, for example, by age (e.g., older than two years) or by project directory.

**Hardware**: All tests were run on an Intel(R) Xeon(R) CPU E-2146G CPU with six cores, 31GiB RAM, and a 1 TB PC601 NVMe SK hynix SSD.

**Software and settings**: OS was openSUSE Leap 15.1 with non-default kernel 4.12.14-lp151.28.91. The old kernel version was used for comparison with *Burrito*, whose *systemtap*-script from 2012 does not run on more recent kernel versions. The benchmark used a dedicated disk to reduce the impact of OS-activity on the results. For the same reason, multi-queue disk access (Bjørling et al., 2016) was enabled. Native command queuing (NCQ) was disabled due to potentially large, don-deterministic delays and vendor-specific implementation (Jose et al., 2008). To make benchmark executions comparable and stable, they were executed at the highest non-turbo CPU-frequency with Hyper-threading disabled. As modern processors often control the frequency themselves, rather than following user requests, cores were kept near their maximum frequency using the `idle=poll` kernel parameter. The CPU scaling driver `intel_pstate` was disabled in favor of the older `acpi-cpufreq` driver to reduce noise, as reported in (Dorn et al., 2017). Each benchmark iteration started with a “cold” cache: disk cache was cleared by reading 1.5 GB of unrelated data from the benchmark drive, and the OS page-cache (Love, 2005, chap. 15) was evicted by re-mounting the respective partitions after each run. The full Grub command line was:

```
idle=poll intel_pstate=disable intel_idle.max_cstate=0
processor.max_cstate=1 scsi_mod.use_blk_mq=1
libata.force=noncq,noncqtrim systemd.unified_cgroup_hierarchy=0
```

The tracing tools were used in the following versions: *shournal* (v2.9), *Burrito* (commit 6630cb2), SPADE (commit 3437fcd), strace (v4.20). For the *kernel copy* the Linux source code v4.19.132 was copied with the *cp* command. During *git checkout* the Linux git tree, hard resetted to v4.19, was checked out to v3.10. Finally, *make* was executed on elfutils v0.176 using `configure && make -j$(nproc)`.

Due to the relatively small overhead of the kernel module backend, 300 repetitions were performed for each command. All other backends (including *fanotify*) were configured for 100 repetitions. In all cases, runtimes were recorded after seven “warmups”. To compensate for the potential long-lasting cache effects, traced and untraced runs were carried out alternating, while, for each execution pair, it was randomly sampled which to run first.

*shournal* was configured to impose no limit on the number of logged read and written files, to store at most ten read scripts ending with the suffix *.sh*, and not exceeding a size of 0.5MiB and to calculate partial file checksums with the default of at most $3 \bullet 256$ bytes. As described previously, events are logged to an intermediate binary file during tracing, while the final storing to the sqlite database happens later. Therefore, that final data transfer was not part of the runtime measurements. We consider this benchmark design reasonable since, after the time-critical recording phase, the collected provenance can be held indefinitely long and is permanently stored using a low-priority background thread.

Event logging was performed on the same disk where the respective files of the benchmark resided, constituting a worst-case.

### 3 Limitations

To our knowledge, *shournal* is currently the only readily available tool to associate shell commands and their used files with low overhead, irrespective of the underlying filesystem. However, in its current implementation (version 3.0), *shournal* still lacks some desirable features. For example, the provenance of binary executables is not tracked. Therefore, changes in binary executables may impair reproducibility. Further, the file event history may have gaps when calculations are distributed over multiple machines. While provenance is recorded on all devices where *shournal* is installed, the reconstruction of the history is more challenging in this case. Both features shall be supported in upcoming versions of *shournal*. Files opened read-writable are currently handled as write-only files. While many workflows clearly separate input- and output files in practice, there are exceptions. Future *shournal* versions shall at least provide a means to flag such files as potential input candidates.

Tracking is currently unavailable when the observed process $A$ instructs the unobserved process $B$ via interprocess communication (IPC) (Kerrisk, 2010, chap. 43.1) to use a file. For shell workflows, this is rather uncommon. However, if it proves to be an issue, future versions of *shournal* shall allow executing $B$ in the provenance context of $A$.

In cases of extremely high file event frequencies of a single process tree, *shournal* may drop some events to avoid slowing down the process at the cost of losing the provenance of individual files. Apart from artificial benchmarks, we observed this rarely for the *fanotify* backend and never for the kernel module backend.

Memory-mapped (Kerrisk, 2010, chap. 49) files are currently only tracked correctly if the underlying file descriptor is closed after (and not before) usage. While this is good practice, it is, technically, not required by Linux. Thus, events evoked by programs that do not properly close the descriptor may be missed.

In the case of the *fanotify* backend, the observation of a process tree ends when the last instance of a dedicated file descriptor, inherited by child processes (Kerrisk, 2010, chap. 2.5), is closed (which happens automatically on exit). If processes do not wait on their children **and** explicitly close this file descriptor, subsequent file events are lost. Further, as process trees are isolated against each other using unshared mount namespaces, file provenance for traced programs which unshare the mount namespace themselves is lost. The kernel module backend does not suffer from these limitations and can be used instead if necessary.

When a dedicated file descriptor is opened during interactive sessions (e.g., via `exec 3> some-file`) and used within separately entered commands, only the last command using it before closing is associated with that file event. The background is that *shournal* only tracks close events when the last instance of a given file descriptor is closed. The same applies, e.g., when the descriptor is sent via *unix domain sockets* (Kerrisk, 2010, chap. 57) to an unobserved process, and the final closing happens there. In practice, however, this should not be an issue.

### 4 Applicability of shournal

To get an intuitive understanding of the applicability of our tool, we provide a simple usage example.

Scientist Sarah creates a new pipeline for her current project. As this work involves a lot of trials and a careful examination of each intermediate result, all of the steps are first carried out on the interactive shell. In the middle of the process, her collaborator asks her for another short but urgent analysis. Sarah does this and returns to the initial project. Unfortunately, she has no idea where exactly she left off. If she is lucky, the last steps are still within the shell's history. This reconstruction, however, involves searching through unrelated commands executed in between. Fortunately, Sarah organizes her work in project directories and can quickly ask *shournal* what she did last in that particular working directory (cwd). By typing `shournal -q -cwd $PWD --history 50`, Sarah can recall the last 50 entered commands, including a brief summary of the read and written files. Sarah then quickly resumes and finishes the analysis. Finally, she wants to automate the steps by writing a shell script.

Initially, she aligned RNA-sequencing reads with a certain accuracy threshold, which resulted in a file `rna.bam`. But which threshold was it exactly? The shell's history shows she tried a good dozen different ones. Since *shournal* was previously configured to capture the provenance of all files written to that project directory, she is able to answer this question very quickly. Executing `shournal -q -wf rna.bam` yields exactly the shell command that led to the creation of this file and unveils the threshold 95. Of note, the `-wf`-(written-file-)argument ensures that this command works irrespective of the filename `rna.bam`, as the file is found in *shournal*'s database by its metadata (size, modification time) and hash. Next, Sarah executes the whole script, and all goes well. The next day she remembers documenting her work and creating an archive of the experiment using, e.g., ReproZip (Chirigati et al., 2016). Unfortunately, the result is suddenly different from the previous run. While this was the beginning of a frustrating process in the past, finding the reasons for this difference is more straightforward with *shournal*. By calling a *diff* on the read files of the two executed commands, it becomes clear that a co-worker modified a common script which is part of Sarah's pipeline. Since Sarah configured *shournal* to store script files (files ending with .sh) within its database, reproducing her previous result takes only minutes. Even in cases where the whole script file itself was not archived, the different metadata (size, modification time) and the hash of a changed read or written file is a strong hint where action is needed. Finally, Sarah does not only work on the shell but also with the workflow manager *Snakemake* (Köster and Rahmann, 2012) for recurring tasks. Using the converter shournal-to-snakemake, the process of transforming the existing pipeline into a *Snakefile* is simplified.
